# High levels of Dorsal transcription factor downregulate, not promote, *snail* expression by regulating enhancer action

**DOI:** 10.1101/2021.01.11.426256

**Authors:** Jihyun Irizarry, James McGehee, Angelike Stathopoulos

## Abstract

In *Drosophila* embryos, genes expressed along the dorsal-ventral axis are responsive to concentration of the Dorsal (Dl) transcription factor, which varies in space; however, levels of this morphogen also build over time. Since expression of high-threshold Dl target genes such as *snail* (*sna*) is supported before Dl levels peak, it is unclear what role increasing levels have if any. Here we investigated action of two enhancers that control *sna* expression in embryos, demonstrating using genome editing that Dl binding sites within one enhancer located promoter proximally, *sna.prox*, can limit the ability of the other distally-located enhancer, *sna.dis*, to increase *sna* levels. In addition, MS2-MCP live imaging was used to study *sna* transcription rate in wildtype, *dl* heterozygote, and a background in which a photo-sensitive degron is fused to Dl (*dl-BLID*). The results demonstrate that, when Dl levels are high, Dl acts through *sna.prox* to limit the activity of *sna.dis* and thereby influence *sna* transcription rate. In contrast, when Dl levels are kept low using *dl-BLID, sna.prox* positively influences *sna* transcription rate. Collectively, our data support the view that Dl’s effect on gene expression changes over time, switching from promoting *sna* expression at low concentration to dampening *sna* expression at high concentration by regulating enhancer interactions. We propose this differential action of the Dl morphogen is likely supported by occupancy of this factor first to high and then low affinity binding sites over time as Dl levels rise to coordinate action of these two co-acting enhancers.

**Significance statement:** A gradient of the maternal transcription factor Dorsal is important for establishing spatial expression of target genes along the dorsal-ventral axis of *Drosophila* embryos. Dorsal levels are also dynamic as nuclear concentration builds in time. Surprisingly, expression of high-threshold target genes such as *snail* is supported before levels peak, raising the question why levels continue to build. Our data support the view that peak Dorsal levels act to preferentially support activity of one enhancer over another to effectively decrease *snail* expression. In addition, while the morphogen Dorsal acts early to support gene expression, later it effectively acts as a damper to limit gene expression. Our results suggest other morphogens also have effects on gene expression that change over time.

## Introduction

Recent studies have shed light on the importance of the timing of morphogen exposure during development and the duration of morphogen input on target gene expression (rev. in 1, 2–4). Despite accumulating data supporting temporal roles for morphogens during development, it remains unclear in most cases whether morphogens directly support multiple actions over time and/or include indirect responses of the underlying gene regulatory networks. In this study, we investigated the temporal action of Dorsal (Dl), an atypical morphogen and transcription factor, which patterns the dorsal-ventral (DV) axis in a concentration-dependent manner. While genes expressed in ventral and lateral regions along the DV axis are responsive to Dl levels (5), we found that some target genes such as *snail* (*sna*), once initiated, no longer require Dl-input to sustain expression (6). Dl becomes dispensable for *sna* expression once the early Dl-target gene *twist* (*twi*) is expressed, demonstrating that the gene network functioning downstream of Dl, an indirect response, takes over late. However, Dl levels also build in time (7) and it remains unclear what role higher levels of Dl support if levels present earlier are sufficient to support gene expression of high-threshold targets such as *sna*.

In particular, we hypothesized that Dl’s effect on gene expression changes over time, depending on its interaction with distinct enhancers driving particular genes. Previous studies have demonstrated that two enhancers regulate *sna* gene expression in the early embryo, one located in a promoter proximal position (*sna.prox*) and the other located distally (*sna.dis*) (8–11). *sna.prox* includes a ∼2 kB region bound by Dl, demonstrated by both chromatin immunoprecipitation (ChIP) and footprinting (9, 12). *sna.dis* also is bound by Dl, identified through ChIP (12, 13). Furthermore, previous studies using large reporter constructs to assay the individual roles of *sna.prox* and *sna.dis* demonstrated that these two enhancers differentially influence levels of *sna* gene expression (8, 14). In particular, deletion of the *sna.prox* enhancer in the context of large reporter constructs results in higher levels of *sna* expression when examined during nuclear cycle (nc) 14 (8, 14), supporting the view that this sequence normally acts as a “damper” to decrease *sna* expression levels during this time. In this study, we investigated how Dl levels impact *sna* expression and, in particular, examined the role of *sna.prox* enhancer over time.

## Results

### *Deletions of* sna *embryonic enhancers from the endogenous* sna *locus lead to changes in* sna *expression levels*

To examine the roles of *sna.prox* and *sna.dis* in the native context, we used Crispr-Cas9 genomic editing to delete sequences associated with each enhancer from the *sna* gene locus (Fig. 1A). When a 1.8kB region encompassing the distal enhancer is deleted (Fig. 1A, *Δdis1.8*), *sna* expression levels are lower than wildtype when qPCR is used to measure total transcript levels at late nc14 (i.e. nc14c, Fig. 1C,F, compare with 1B). Conversely, when a 4.4kB segment encompassing the proximal enhancer (Fig. 1A, *Δprox4.4*) is removed, *sna* expression level is higher than wildtype (Fig. 1D,F). Similar results were obtained with analysis of enhancer deletions in the context of large reporter constructs encompassing ∼25 kB surrounding *sna* locus, suggesting that the proximal enhancer drives weak *sna* expression levels, whereas the distal enhancer supports strong *sna* expression levels (8, 14). Previous studies had suggested that coordination of proximal and distal enhancers is required to support correct *sna* expression levels (8), therefore we used genomic editing to create mutant lines with different *sna.prox* enhancer deletions to identify the minimal portion able to properly control *sna* levels of expression.

**Figure 1.**
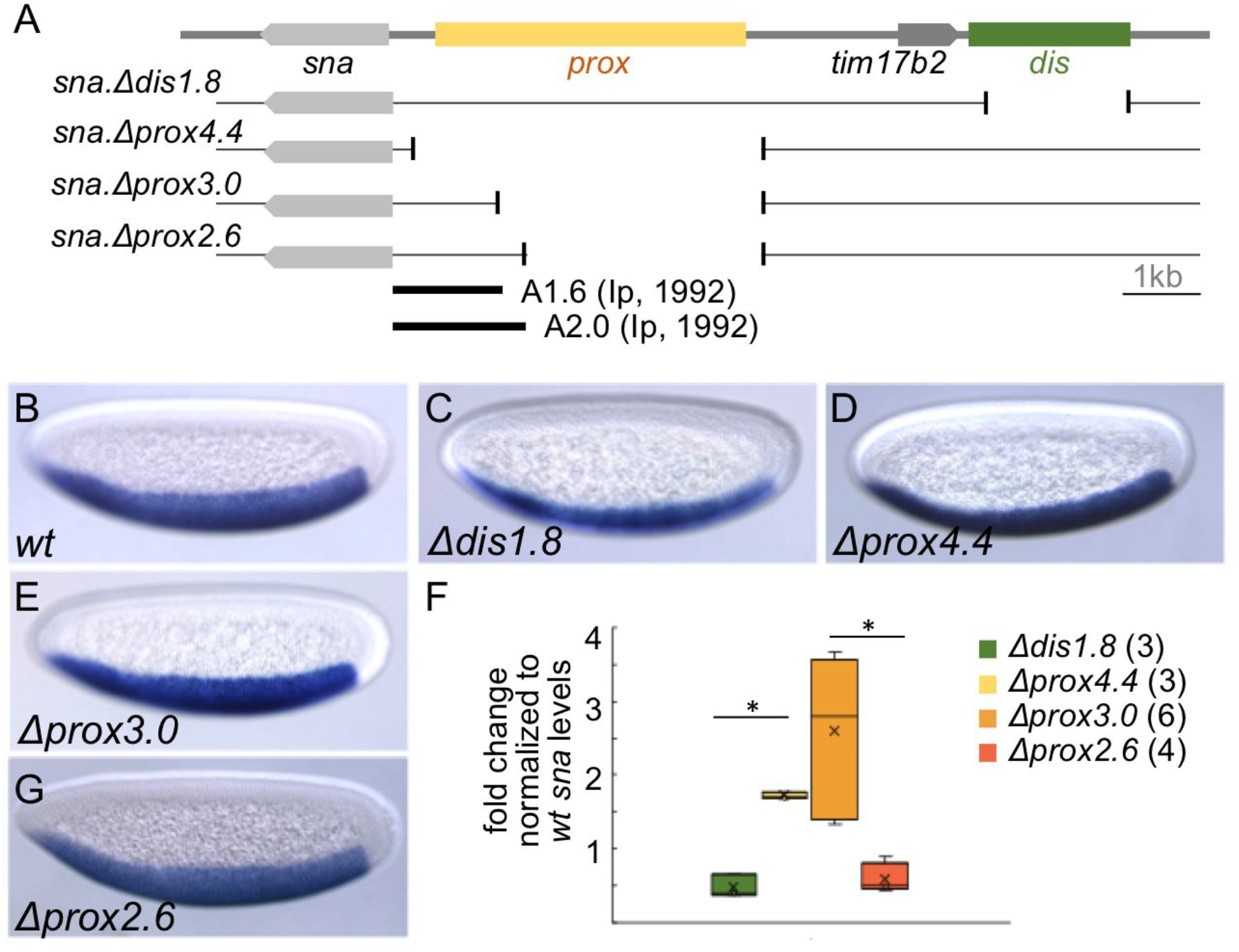
Embryos with various deletions of *sna* distal or proximal enhancers fail to support endogenous *sna* expression levels. (A) Schematic of enhancer deletions, proximal (*prox*, yellow) or distal (*dis*, green), by CRISPR-Cas9 system. Two regions in the proximal enhancer identified to support incomplete or full *sna* expression, A1.6 or A2.0, respectively (9). (B-E,G) *sna* expression in *Drosophila* nc14d embryos by *in situ* hybridization using *sna* riboprobes. Lateral views of embryos are shown, with anterior to the left and dorsal side up. (F) Normalized *sna* transcript levels in embryos at nc14c for indicated deletion mutations relative to wildtype determined using qPCR. Sample sizes are indicated next to the genotype. Asterisk: p<0.01 by Student’s t-test. For this and all subsequent figures, box plots show median (center line) and 25th or 75th percentiles (box limits). Whiskers represent 1.5 times the interquartile range from the 25th and 75th percentiles.

Two mutant lines were generated with different sized deletions of the proximal enhancer region. These deletions were positioned using information obtained from previous reporter analysis that had highlighted important sequences (Fig. 1A). A 2.0kB portion of the proximal enhancer (i.e. A2.0) was shown to support expression similar to endogenous *sna* expression; whereas, a 1.6kB portion of the proximal enhancer (i.e. A1.6) is compromised, supporting only partial expression (9). Using genome editing to modify the endogenous *sna* locus, the first deletion, encompassing a 2.6kB distal sequence (Fig. 1A, *Δprox2.6*) leaves the full, minimally-defined proximal enhancer intact such that full ventral expression remains (i.e. A2.0, Fig. 1A) (9). In *Δprox2.6* embryos, while expression is normal in terms of spatial domain (Fig. 1G), *sna* levels are slightly lower than wildtype (Fig. 1F). In contrast, when a larger 3.0kB region is deleted using genome editing (Fig. 1A, *Δprox3.0*), with a breakpoint that is similar to the minimally-active A1.6 reporter construct (9), *sna* levels are variable and higher than wildtype (Fig. 1F). It was unexpected that embryos that contain a partially active proximal enhancer sequence (i.e. *Δprox3.0*) are associated with higher *sna* gene expression.

### *Partial deletions of the* sna.prox *enhancer lead to an increase in* sna *levels due to uncontrolled* sna.dis enhancer *activity*

Together with the observation that the distal enhancer supports high *sna* expression (8, 14), we hypothesized that the higher *sna* expression found to be associated with *Δprox3.0* relates to an overactive distal enhancer. To test this idea, the distal enhancer was removed from the *Δprox3.0* background, creating double mutants (i.e. *Δprox3.0 Δdis1.8*, Fig. 2A). In *Δprox3.0 Δdis1.8* double mutant embryos, the *sna* levels are decreased relative to wildtype, and similar to the *sna* levels detected in *Δdis1.8* single mutant embryos (Fig. 2B, C, compared with Fig. 1F). This result suggests that the high *sna* levels associated with the *Δprox3.0* mutant background relates to a hyper-active distal enhancer and, additionally, that input from the proximal enhancer is pivotal for regulation of *sna* expression levels.

**Figure 2.**
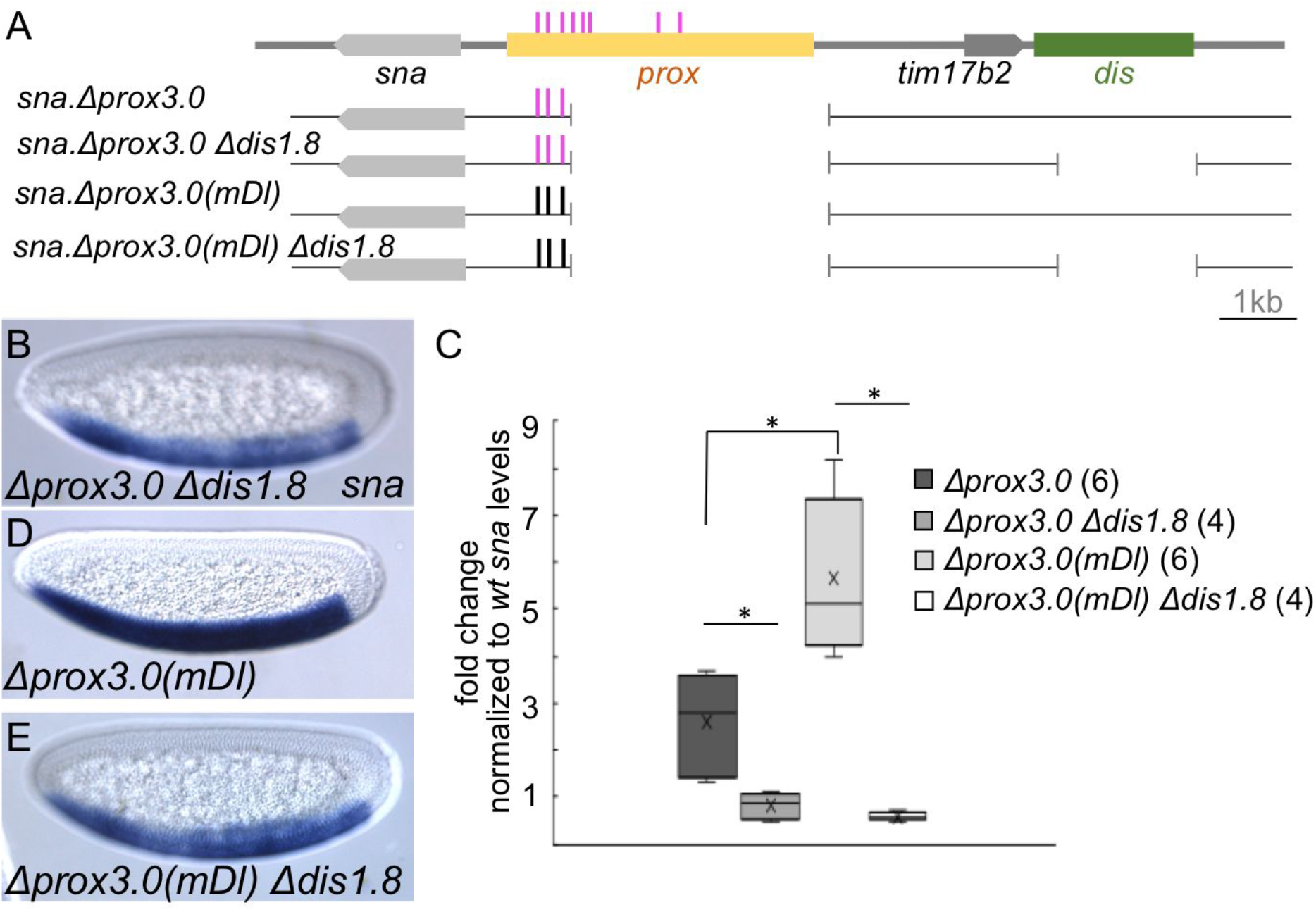
Mutating Dl binding sites in *sna.Δprox3.0* increases *sna* expression level in a distal enhancer-dependent manner. (A) Schematic of deletions in the proximal enhancer with/without the distal enhancer deletion. Dark pink bar: putative high-affinity Dl binding sites in the proximal enhancer identified by JASPAR (http://jaspar.genereg.net). Black bar: mutated putative high-affinity Dl binding sites. (B,D,E) *sna* expression in nc14d embryos by *in situ* hybridization using *sna* riboprobes. Lateral views of embryos are shown, with anterior to the left and dorsal side up. (C) Normalized *sna* transcript levels in embryos at nc14c for indicated deletion mutations relative to wildtype determined using qPCR. Sample sizes are indicated next to the genotype. Asterisk: p<0.01 by Student’s t-test

To further characterize the proximal enhancer’s mechanism of action, we mutated Dl binding sites in the *Δprox3.0* background that we hypothesized contains a partially active proximal enhancer; since mutating these sites in the context of reporter assays was associated with reduction of proximal enhancer activity (9). When three Dl binding sites that match the JASPAR consensus binding site (http://jaspar.genereg.net), likely of high affinity, are mutated in *Δprox3.0* [i.e. *Δprox3.0(mDl)*], Figs. 2A and S1), the resulting *sna* levels are higher than *Δprox3.0* (Fig. 2C; Fig. 2D, compare with 1E). To test whether the increased *sna* levels associated with *Δprox3.0(mDl)* also relate to a hyper-active distal enhancer, the distal enhancer was additionally deleted from this background [i.e. *Δprox3.0(mDl) Δdis1.8*, Fig. 2A]. In embryos containing this double deletion, *sna* levels are lower than wildtype, but similar to levels detected in *Δdis1.8* embryos (Fig. 2C; Fig. 2E, compare with 1E). This result demonstrates that the increased expression associated with mutation of Dl sites within the proximal enhancer stems from increased action of the distal enhancer. Taken together, these results and those in the previous section support the view that a functional proximal enhancer limits activity of the distal enhancer, thereby supporting the physiologically appropriate levels of *sna* expression.

### *High Dl concentration leads to a decrease in* sna *levels by limiting* sna.dis *enhancer activity*

Previous studies have used the MS2-MCP.GFP system to monitor *sna* transcription from large reporter constructs in which the *sna* gene has been replaced by the MS2.yellow dual-reporter sequence and examined the relative roles of *sna.prox* and *sna.dis* in supporting expression over time (14) (Fig. S2A). These studies found that *sna* transcription decreases during the course of nc14 (14), and were independently confirmed using our imaging settings (Fig. S2B). On the other hand, Dl concentrations continuously increase during nc14 during this same period (7), demonstrating an anti-correlation exists between Dl and *sna* levels during this stage. This is unexpected as Dl is thought to be a positive input to *sna* and that, moreover, high levels of Dl are required to support *sna*. We hypothesized that at nc14, Dl’s effect on *sna* expression changes, possibly negatively impacting *sna* levels.

To further investigate this idea that Dl negatively impacts *sna, sna* levels were measured by monitoring *sna* nascent transcription in embryos containing reduced Dl levels (i.e. *dl*^*4*^ heterozygotes) to determine whether transcription is increased. The *sna* MS2 reporter construct was introduced into a *dl*^*4*^*/+* background and the mean spot size (i.e. mean number of pixels per MS2-MCP.GFP spot), which is correlated to the rate of transcription (15), was examined in early embryos. Initially, at early nc14, the mean spot size is lower in *dl*^*4*^ heterozygotes than wildtype (Fig. 3B t=4; Fig. 3C and Movie S1). However, as *dl*^*4*^ heterozygous embryos age, the mean spot size increases relative to wildtype (Fig. 3B t=44; Fig. 3C and Movie S1). We hypothesized that this trend relates to decreased proximal enhancer activity as a result of limiting Dl.

**Figure 3.**
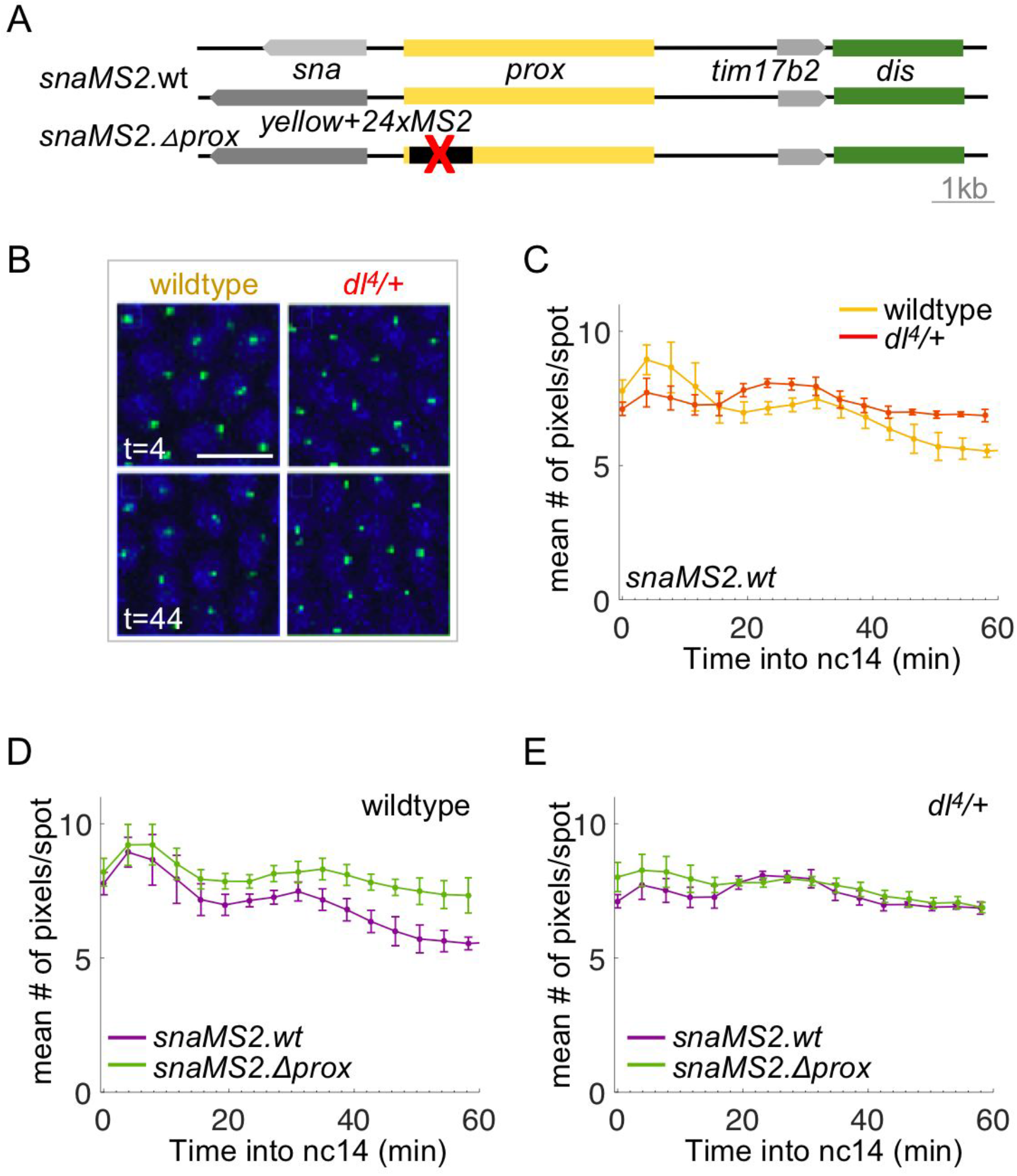
At high Dl levels, the proximal enhancer limits the distal enhancer activity, consequently decreasing *sna* expression levels. (A) Schematic of reporter constructs used to assay *sna* transcriptional activity by monitoring MS2-MCP.GFP (14). (B) MCP.GFP signal associated with *snaMS2.wt* (green dots) driven by both enhancers (“prox+dis acting”) in wildtype (left column) or *dl*^*4*^*/+* (right column) embryos at time point 4 min (t=4) or 44 min (t=44) after entering nc14. Nuclei are visualized by H2A-BFP (blue). Scale bar: 10μm. (C) Quantitative analysis of mean number of pixels per MCP.GFP spot associated with *snaMS2.wt* in wildtype (yellow, high Dl levels) and *dl*^*4*^*/+* (red, intermediate Dl levels) embryos. (D,E) Quantitative analysis of mean number of pixels per MCP.GFP spot associated with *snaMS2.Δprox* (green) in wildtype (D) or *dl*^*4*^*/+* heterozygotes (E). The mean number of pixels per MCP.GFP spot associated with *snaMS2.wt* (purple) in wildtype or *dl*^*4*^*/+* heterozygotes used in C are replotted for direct comparison to mean number of pixels per MCP.GFP spot associated with *snaMS2.Δprox*. Error bars represent standard error of the mean. For each condition, movies were obtained for three individual embryos and the data averaged. For the individual traces, see Figure S4.

To test this idea, we measured the mean spot size associated with *snaMS2.Δprox*, in which only the distal enhancer is active, in a wildtype or *dl*^*4*^ heterozygous background, and these levels were compared to the levels supported by both enhancers (i.e. *snaMS2.wt*). In wildtype embryos, in which Dl levels are high, the mean spot size associated with *snaMS2.Δprox* is higher than *snaMS2.wt;* most clearly demonstrated at late nc14 (Figs. 3D, S4A, and Movie S2). This suggests that input from the proximal enhancer normally dampens *sna* levels. In contrast, in *dl*^*4*^ heterozygotes at this same stage, no difference is observed in the mean spot sizes of *snaMS2.wt* or *snaMS2.Δprox* (Figs. 3E, S4B, and Movie S2). These results demonstrate that reducing Dl levels is equivalent to greatly reducing *sna.prox* activity, and also suggest that the Dl levels present in a *dl*^*4*^/+ background are lower than necessary to support proximal enhancer activity.

### *Decreased levels of Dl support cooperative action of* sna.prox *and* sna.dis *enhancers*

To further investigate the idea that Dl and *sna* levels anti-correlate, we used the MS2-MCP.mCherry system (16) to examine *sna* levels *in vivo* that result in real-time from controlled modulation of Dl levels accomplished using a previously defined optogenetic approach to degrade Dl. In brief, this optogenetic approach supports degradation of Dl with fine-scale temporal resolution by fusing a sequence encoding a Blue Light-Inducible Degron (BLID) to the C-terminus of Dl (Dl-BLID) at the endogenous locus (6, 17). In the dark, the degron domain is inaccessible and Dl-BLID protein is present and functional; whereas, upon exposure to blue light, the degron is exposed, resulting in Dl-BLID protein degradation in less than 5 min (17). In this previous study, we used MCP.GFP to monitor nascent transcripts from *sna.MS2* reporters upon blue light application, but we were unable to monitor the ground state (before illumination with blue laser) because excitation of GFP also supported activation of the photosensitive degron. To address this issue, we used MCP.mCherry to monitor *sna.MS2* stem loop production. This allows us to monitor *sna* transcriptional activity in nc14c before blue laser illumination is applied, which serves as a reference/ground state, and also to monitor subsequent effects after blue light illumination, until nc14d (Fig. 4A). As a control, wildtype embryos devoid of any *dl-BLID* fusion were illuminated with a blue laser while detecting MCP.mCherry signal associated with *snaMS2.wt* (Fig. 4A). Only a slight decrease in mean spot size was observed between embryos with 5 min blue laser illumination relative to embryos kept in dark (Figs. 4C, S5A, S5D, and Movie S5).

**Figure 4.**
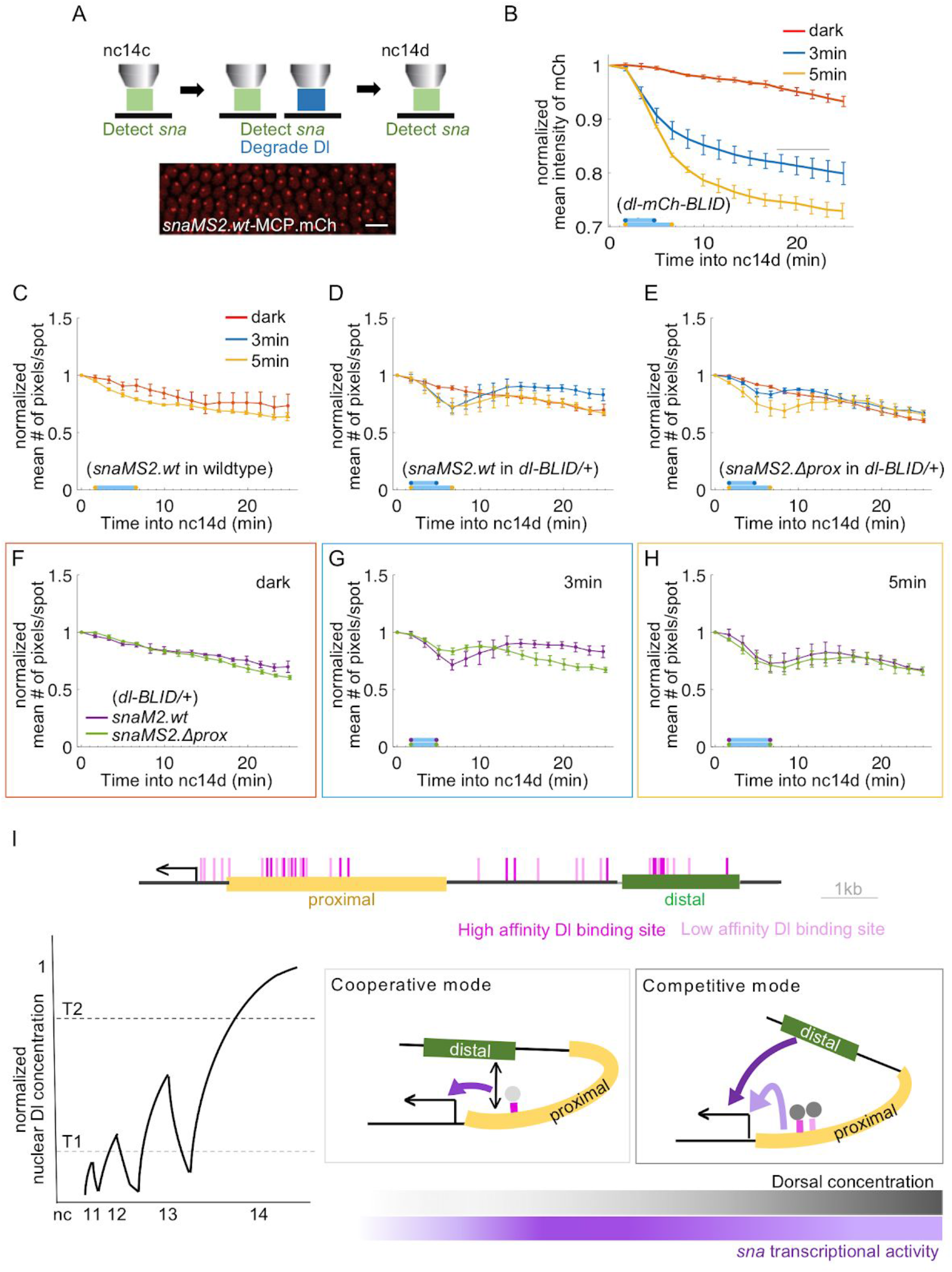
At low Dl levels, the distal enhancer activity is dependent on the proximal enhancer to support endogenous *sna* expression levels. (A) Schematic of *sna* MS2-MCP.mCherry system combined with the Dl-BLID degradation system at late nc14. MCP.mCherry signal associated with *sna* MS2 (red dots) driven by both enhancers (*snaMS2.wt*) in wildtype embryos are shown. Scale bar: 10μm. (B) Quantitative analysis of normalized mean intensity of mCherry fused to Dl-BLID proteins [*dl-mCh-BLID*; (6). *dl-mCh-BLID* embryos were kept in dark or illuminated by a confocal blue laser for 3 or 5 min. Mean intensity of mCherry associated with Dl-BLID is measured in each time frame, and the values are normalized to the initial value detected in the first frame. For the individual traces, see Figure S5. (C,D) Quantitative analysis of normalized mean number of pixels per MCP.mCherry spot associated with *snaMS2.wt* in wildtype (C) or *dl-BLID/+* embryos (D). (E) Quantitative analysis of normalized mean number of pixels per MCP.mCherry spot associated with *snaMS2.Δprox* in *dl-BLID/+* embryos. Embryos are kept in dark (red) or illuminated with a confocal blue laser for 3 min (blue) or 5 min (yellow). For the individual traces, see Figure S6. (F-H) Normalized mean number of pixels per MCP.mCherry spot associated with *snaMS2.wt* (purple) or *snaMS2.Δprox* (green) in *dl-BLID/+* embryos kept in dark (F) or with 3 min (G) or 5 min (H) blue laser illumination are replotted with the date used in D and E. Mean spot size of MCP.mCherry associated with *sna* MS2 are measured in each time frame, and the values are normalized to the initial value detected in the first frame. Error bars represent standard error of the mean. For each condition, movies were obtained for three individual embryos and the data averaged. (I) Schematic showing high (dark pink) or low affinity (light pink) Dl binding sites predicted by JASPAR (http://jaspar.genereg.net) at *sna* locus (top). A conceptual representation of Dl (bottom left) from nc11 to nc14 (7). The low threshold (T1: threshold 1) supports cooperative action of enhancers, and the high threshold (T2: threshold 2) supports enhancer competition. Model showing two modes of coordination between the proximal and distal enhancers, dependent on the concentration of Dl, to support the correct level of *sna* expression (bottom right).

To confirm that different levels of Dl degradation can be achieved with blue laser illumination of embryos for varying lengths of time, flies with a mCherry-BLID domain fused to the C-terminus of *dl* at the endogenous locus (i.e. *dl-mCh-BLID*) were utilized so that Dl protein could be tracked (6). This mutant line allows detection of Dl levels by monitoring mCherry fluorescent signal in the presence or absence (dark) of blue laser illumination. To generate low or very low Dl levels, *dl-mCh-BLID* embryos were illuminated for 3 min and 5 min, respectively. As expected, *dl-mCh-BLID* embryos with 3 min blue laser illumination exhibited lower Dl levels compared to embryos without illumination, whereas *dl-mCh-BLID* embryo with 5 min blue laser illumination exhibited even lower levels of Dl (Figs. 4B, S5, and Movies S3, S4). Since increasing illumination results in successively lower levels, we refer to the levels reached after 3 min illumination as intermediate and the levels reached after 5 min illumination as low.

To assay whether levels of Dl differentially impact *sna* transcription, *dl-BLID* heterozygous embryos were illuminated for 3 min or 5 min to degrade Dl to intermediate or low levels, respectively, while detecting MCP.mCherry signal associated with *snaMS2.wt* (“prox and dis acting”). *dl-BLID* heterozygotes (*dl-BLID*/+) were utilized for *sna.MS2* assays because the *dl-BLID* homozygous embryos (*dl-BLID/dl-BLID*) exhibit only partial *sna* expression. When *dl-BLID* heterozygous embryos containing the *snaMS2.wt* reporter are illuminated, the mean spot size associated with the MS2-MCP.mCherry signal exhibits an incremental decrease during the period of illumination (Figs. 4D, S6B, and S6E), suggesting *sna* transcription decreases as Dl levels decrease. However, following 5 min illumination, the mean spot size recovers to the level present before the illumination (Fig. 4D and Movie S7). In contrast, in the case of only a 3 min illumination, levels rebound to a state higher than that observed with the 5 min illumination or with embryos kept in dark (Figs. 4D, S6A, S6D, and Movie S7 compare with Movie S6). To test whether this rebound requires action of the *sna.prox* enhancer, we assayed the effect of reducing Dl on *snaMS2.Δprox* (i.e. “only dis acting”) reporter output. In this case, the mean spot size continuously decreases after light illumination and no rebound above control (dark) is observed (Figs. 4E, S6C, S6F, and Movies S8, S9). This ability of the *snaMS2.wt* reporter to achieve a higher level of expression after blue laser illumination is (i) *sna.prox* enhancer-dependent (lost in *snaMS2.Δprox*) and (ii) Dl-levels dependent (only associated with 3 min not 5 min illumination). These results also suggest that when low levels of Dl are present, the *sna.prox* enhancer promotes *sna* transcription; in contrast to the enhancer deletion analysis, which supports the view that the proximal enhancer acts to dampen *sna* levels (e.g. Figs. 3D and S1).

We propose that the role of the *sna.prox* enhancer can change in a manner dependent on Dl levels. In embryos illuminated for 3 min, the mean spot size associated with *snaMS2.wt* is larger than that associated with *snaMS2.Δprox* (Fig. 4G). Whereas Illumination of embryos for 5 min (i.e. more Dl-degradation resulting in low Dl levels) results in no difference in the mean spot size associated with *snaMS2.wt* or *snaMS2.Δprox* (Fig. 4H), suggesting that a critical level of Dl is necessary to support proximal enhancer action. Taken together, when Dl levels are low but not high enough to support the proximal action, the distal enhancer acts alone to drive weak *sna* levels; when Dl levels are a bit higher but not high enough to support “damper” action by the proximal, the distal and proximal enhancers work cooperatively (Fig. 4I). Under wildtype conditions with normal Dl levels present, the role of *sna.prox* enhancer changes to support dampening of *sna* levels. The observation that Dl levels build in time has implications for a changing role of the *sna.prox* enhancer over time.

## Discussion

Collectively, our results suggest that initially the proximal and distal enhancers cooperate under conditions of lower Dl but then as Dl levels rise, the proximal enhancer becomes more potent, gaining influence to the promoter over the distal enhancer and subsequently leading to dampening of levels (Fig. 4J-L). While *sna* is considered to be a high-threshold Dl target gene, meaning the highest levels of Dl are required to support its expression compared to other Dl target genes (18), our results suggest that Dl may exhibit a changing effect on *sna* expression due to its action at the *sna.prox* enhancer. It is possible that Dl’s changing effect over time relates to its ability to differentially occupy binding sites of varying affinity within each enhancer (Fig. 4I). Earlier when lower levels of Dl are present, high affinity sites associated within one or both enhancers are occupied (cooperative mode, Fig. 4I); subsequently, low affinity binding sites, which happen to be enriched at the proximal enhancer, are occupied later when peak Dl levels are achieved. Occupancy of low affinity binding sites within the *sna.prox* enhancer may allow this particular enhancer to compete better later and thus dampen gene expression at the later stage (competitive mode, Fig. 4I).

In this study, our data demonstrate that in addition to regulating the spatial domain of *sna* expression, the morphogen Dl additionally acts to regulate levels of expression of this target gene. It is possible that other morphogens also change their perceived effect on gene expression over time, such as changing from promoting gene expression to dampening gene expression. We propose that morphogens commonly provide feedback to target genes, after specifying spatial domains of expression, to help keep target gene expression robust.

Furthemore, the *sna* gene may have acquired two regulatory modules that differentially respond to morphogen Dl concentration as a mechanism to buffer genetic and environmental fluctuations in Dl levels. This idea had been proposed previously as deletion of *sna.prox* in the context of rescue transgenes also was associated with sensitivity to growth at high temperature (8, 11). Here we show that deletion of either enhancer supports viability, but deletion of the *sna.prox* enhancer, specifically, resulted in increased variability in *sna* levels (Fig. 1F). Our results that Dl input to *sna.prox* supports dampening of gene expression, provides a molecular explanation for how *sna* expression can be kept in check despite variability in transcription factor inputs through action of the *sna.prox* enhancer. Animals may have developed such safety mechanisms, such as multiple co-acting enhancers, to overcome the challenges posed by fluctuating morphogen gradients that can result from genetic and environmental perturbations (11, 19). Dual-input by morphogens may increase precision of target gene outputs.

## Materials and methods

### Fly stocks/husbandry and crosses

All flies were kept at 25°C, except for flies bearing the *dl-BLID* allele that were kept at 18°C. *yw* is used for wildtype, unless otherwise noted. To test *sna* transcriptional activity in the *dl*^*4*^ heterozygous background (#7096, Bloomington Drosophila Stock Center), *dl*^*4*^/CyO flies were crossed with *MCP.GFP, mCherry-PCP, His2Av-eBFP2* to generate *dl*^*4*^/CyO; *MCP.GFP, mCherry-PCP, His2Av-eBFP2*. Virgin flies with *MCP.GFP, mCherry-PCP, His2Av-eBFP2* or MCP.mCherry (from Hernan Garcia, UC Berkeley, US; 16) in wildtype, *dl*^*4*^ heterozygous or *dl-BLID* heterozygous backgrounds were crossed with males having both proximal and distal enhancers (WT, *snaMS2.wt*), a proximal deletion (NoPrimary, *snaMS2.Δprox*), or a distal deletion (NoShadow, *snaMS2.Δdis*) (from Michael Levine, Princeton University, US; 14) for live imaging to detect *sna* MS2 stem loops. The *dl-BLID* line was generated in a previous study by integrating BLID domain at the C-terminus of Dl at the endogenous locus using CRISPR/Cas9 (6).

### CRISPR-Cas9 Mediated Genome Modification

To target deletions of various sizes to the distal part of the proximal enhancer [i.e. *sna.Δprox4.4, sna.Δprox3.0, sna.Δprox3.0(mDl)*] or distal enhancer (i.e. *sna.Δdis1.8*), a transgenic line was generated expressing two guide RNAs (gRNAs) targeting the flanking region described as the proximal enhancer or distal enhancer (8). First, the unique PAM recognition sites were identified flanking this region using the flyCRISPR optimal target finder (https://flycrispr.org/target-finder). To delete the full proximal enhancer, two flanking regions that are located outside of the identified proximal enhancer (8) were targeted due to limited availability of PAM sequences, and subsequently led to the generation of a 4.4kB deletion. These two sites were cloned into the plasmid pCFD4-U6:1_U6:3tandemgRNAs (Addgene plasmid #49411). The plasmid was injected into *y2cho2v1; P {nos-phiC31\int.NLS}6X; attP2 (III)* (NIG-Fly #TBX-0003), resulting in phiC31-mediated site-integration at landing site attP2 (Chr. III) (20). Integration in the genome at this position was confirmed by PCR/sequencing.

To generate the proximal deletion lines [*sna.Δprox3.0, sna.Δprox2.6 sna.Δprox3.0(mDl)*] and the full distal deletion (*sna.Δdis2.0*), Homology-Directed Repair (HDR)/CRISPR-Cas9 was used (21). A donor construct was generated using pHD-DsRed vector (Addgene plasmid #51434). An ∼1kb 5’ or 3’ homology arm, either upstream or downstream of the region, was cloned with SmaI/NheI or AscI/XhoI, respectively.

*y2cho2v1;sp/CyO;P {nos-Cas9,y+,v +} 2A* (NIG-Fly #Cas-0004) virgin flies were crossed with gRNA transgenic male flies. Embryos were collected and injected with 300 ng/μl of the donor vector. The deletion lines were screened by DsRed expression in adult fly eyes and sequencing. The DsRed marker was removed by crossing with a Cre expressing fly line (y[1] w[67c23] P{y[+mDint2]=Crey}1b; D[∗]/TM3, Sb[1], BDSC #851). Crispr/Cas9 generated deletions/mutations were confirmed by PCRing the region of interest and subsequently sequencing. The sequences after Cre-mediated DsRed marker excision are provided below; uppercase indicates genomic sequence, and lowercase indicates loxP remnant sequence.

#### >sna.Δprox3.0

CGACAAAGGATGTGACTCAGcggccgcggacatatgcacacctgcgatcgtagtgccccaactggggtaacctttgagttctctcagttgggggcgtagataacttcgtataatgtatgctatacgaagttatagaagagcactagtATGACCCACCAGGTA GGATG

#### >sna.Δprox2.6

AGTCGACAAAGGATGTGACTCAGcggccgcggacatatgcacacctgcgatcgtagtgccccaactggggtaacctttgagttctctcagttgggggcgtagataacttcgtataatgtatgctatacgaagttatagaagagcactagtTCTCAGCAAAAATTGACAAGAACAACAA

#### >sna.Δdis2.0

TAAATTCCAACATTTTGCTGcatatgcacacctgcgatcataacttcgtataatgtatgctatacgaagttatagaagagcactagTAAATCCGTGTTAAATTGTT

To introduce mutations into the predicted Dl binding sites of *sna.Δprox3.0* (see Figure 2A), first all three Dl consensus sequence were replaced with the complementary base (i.e. A>T or G>C), and 1kb of the proximal enhancer sequence including all of the mutated Dl consensus sequence was synthesized and inserted into pUC57 (GenScript). This plasmid was used as a DNA template to generate the homology right arm for HDR donor construct.

#### >sna.Δprox3.0(mDl)

(uppercase: genomic sequence, lowercase: mutated Dl consensus sequences)

ATGACCCACCAGGTAGGATGTGAGGACATAatcttttggggAGCCAGTTTTTCCACTCGTCGTGGCTTGTTTTGCTTGAGTTTCGCTGACTGCGTAATTGGATAAGATGGGAAATTACTTTAAATCCTTCGCTGATCCACATCCGGACATTCGTCGAAGGAAAATCCATTGCAGGGAAATACGAAATGGAAATGCGGCTGGGTTATTGGCTCGACATTTCCCATCTTCCCTCACGCCATTGGTTGCAGGATCGCGGGGAAaaccttaaggcgGCTGGAATTTTTTGTCACCTCTTGGGTTTATCAAAACTTTTGGGTTTGCTATcctaaaaaaggtATTTTACC

To generate sna.Δdis1.8 or *sna.Δprox4.4*, non-homologous end joining (NHEJ)/CRISPR-Cas9 was used. y2cho2v1;sp/CyO;P {nos-Cas9,y+,v +} 2A (NIG-Fly #Cas-0004) virgin flies were crossed with gRNA transgenic male flies. The mutant lines with these deletions were confirmed by PCR/sequencing.

#### >sna.Δdis1.8

(uppercase: genomic sequence, lowercase: sequence added after NHEJ) TGAACTTGTTGTGAACTCAGtaccGTCTAATGGCCAGAACACCG

#### >sna.Δprox4.4

TCAAAATCAAAGGGCAGGCCagagtcgacaaaggatgtgacgcAGGGTGCGCCTGCGTCTGCTCTT CC

### In situ *hybridization and imaging*

Embryos were collected for 2hrs, aged at 25°C to give 2-4 hr old embryos, fixed and stained by *in situ* hybridization using standard protocols. Antisense *sna* RNA probes were labeled with digoxigenin. For fluorescent in situ hybridization, sheep anti-digoxigenin (Life Technology PA185378) was used with Alexa conjugate secondaries (1:400; Thermo Fisher). For alkaline phosphatase (AP) staining samples, a Zeiss Axioxam 506 microscope was used, and images were taken with a 20x objective.

### RNA extraction from single embryos and qPCR

Embryos were collected for 1hr and aged for 2hrs at 25°C. They were hand dechorionated and immersed in halocarbon 27 oil to determine the stage of the embryos. When the embryos reached nc14c, in which 75% of cellularization has been completed, the individual embryo was collected in a 1.5ml microcentrifuge tube for RNA extraction using a standard Trizol protocol (ThermoFisher), followed by DNase I treatment (NEB), and cDNA synthesis using SuperScript III First-Strand Synthesis System kit (ThermoFisher). qPCR was performed on cDNA using SYBR Green I Master Mix (Roche) on a StepOnePlus Real-Time PCR System (Applied Biosciences).

### Confocal blue laser illumination

*dl-mCherry-BLID* or *dl-BLID* heterozygous embryos were collected for 1hr and aged for 4hr at 18°C. Embryos were hand dechorionated, and placed onto a heptane glued slide. The specimens were immersed in water and staged using a bright field microscope. Once an embryo reached mid nc14 (i.e. nc14c) determined by the cellularization front having reached 50% completion (∼40 min after entering nc14), the specimen was covered with a cover slip. To degrade Dl, the embryo was illuminated with a Zeiss LSM 800 488nm confocal blue laser with 20% laser power using a 40x water immersion objective (see Fig. 4A).

### Live imaging

To test that *sna* transcriptional activity changes in response to Dl levels, the MS2-MCP system was used. In wildtype (high levels of Dl) and *dl* heterozygote (*dl*^*4*^/+, intermediate level of Dl) embryos, nuclei were marked by imaging H2A-BFP [i.e. His2Av.eBFP2; (22)] with 405nm at 1.5% laser power, and MCP.GFP associated with *sna* MS2 were detected with 488nm at 2.4% laser power. For *dl-mCherry-BLID* or *dl-BLID* heterozygotes, mCherry associated with Dl-BLID or MCP.mCherry associated with *sna* MS2 constructs (16) were detected with 561nm at 5% laser power before, during, and after blue laser illumination (see Fig. 4A). All images taken contained 20 Z-planes separated by 2.0µm.

### MS2 Foci detection and Quantification

MS2 foci were detected and quantified similar to previously described methods (6). Briefly, a gaussian filter was used to smooth the image and background was removed using Top-hat filtering. A threshold was applied to detect MS2 foci. Foci consisting of three or less pixels were removed as often these were not associated with real signal, as confirmed manually, and appeared outside of nuclei or within nuclei already containing a clear focus. For each focus detected, the number of pixels were counted, and the mean intensity of each focus was determined. The number of foci, the mean area (or mean number of pixels per focus), and the average of the mean intensities was computed at every time point for each individual embryo and plotted. Traces were normalized to the first time point before blue laser illumination to account for variability. Error bars were calculated as the standard error of the mean. Codes are deposited at Github under accession number (TBD).

### Dl-mCherry-BLID Quantification

Dl-mCherry-BLID was quantified by making a single z- and time-projection and then drawing a rectangular ROI on the image of the embryo. For each time point, the intensity of each pixel was averaged together for the entire ROI. The same sized ROI was used for all time points of every embryo measured. For each embryo, the ROI was positioned centrally to capture the most signal in the embryo. The size of the ROI was determined beforehand to insure it fit within all embryos measured. Traces were normalized to the first time point before blue laser illumination to account for variability in starting amounts of fluorescence, likely due to variability in Dl concentration. Individual and unnormalized traces are shown in the supplement (Fig. S5). Codes are deposited at Github under accession number (TBD).

## Acknowledgments

We are grateful to Mike Levine for providing the *sna*.MS2, and MCP.GFP fly stocks, Hernan Garcia for providing the MCP.mCherry fly stocks and Leslie Dunipace for comments on the manuscript. This study was supported by funding from National Institutes of Health grant R35GM118146 to A.S.

## Author Contributions

A.S. and J.I. conceived the project and planned the experimental approach. A.S. directed the project. J.I. performed all experiments, and J.M. performed all quantitative analysis of imaging data. Data were analyzed by J.I., J.M., and A.S. The manuscript was written by J.I. and A.S. with edits provided by J.M.

