## Supplemental Information for "High levels of Dorsal transcription factor downregulate, not promote, *snail* expression by regulating enhancer action"

**Supplementary Information for**

High Dorsal levels downregulate, not promote, ventral gene expression in *Drosophila* embryos by regulating enhancer action.

Jihyun Irizarry[1], James McGehee[1], and Angelike Stathopoulos[1,*]

[1]California Institute of Technology, Division of Biology and Biological Engineering, 1200 East California Blvd., Pasadena, CA 91125

**This PDF file includes:**

Figures S1 to S6

Tables S1

Legends for Movies S1 to S9

Legends for Dataset S1

SI References

**Other supplementary materials for this manuscript include the following:**

Movies S1 to S9

A

```
                                   ATCTTTTGGGGG
atgacccaccaggtaggatgtgaggacataTAGAAAACCCCCagccagtttttccactcgtcgtggcttgttttgctt

gagtttcgctgactgcgtaattggataagatgggaaattactttaaatccttcgctgatccacatccggacattcgtc

gaaggaaaatccattgcagggaaatacgaaatggaaatgcggctgggttattggctcgacatttcccatcttccctca
                                AACCTTAAGGCG
cgccattggttgcaggatcgcggggaaTTGGAATTCCGCgctggaatttttgtcacctcttgggtttatcaaaactt
              CCTAAAAAAGGT
ttgggtttgctatGGATTTTTTCCAatttta
```

**Fig. S1. DI binding sites in , and scheme to generate mutants with mutated DI binding sites. a 400 bp segment of the proximal enhancer, and mutations introduced to generate *sna.Δprox3.0(mDI)*.** (A) Predicted DI binding sites identified by JASPER (http://jaspar.genereg.net) were indicated in bold font. The putative DI binding sites were mutated by switching the sequences to complementary sequences (red). Sequences underlined in blue show a subset of DI binding sites identified by DNase I footprint assay (9).

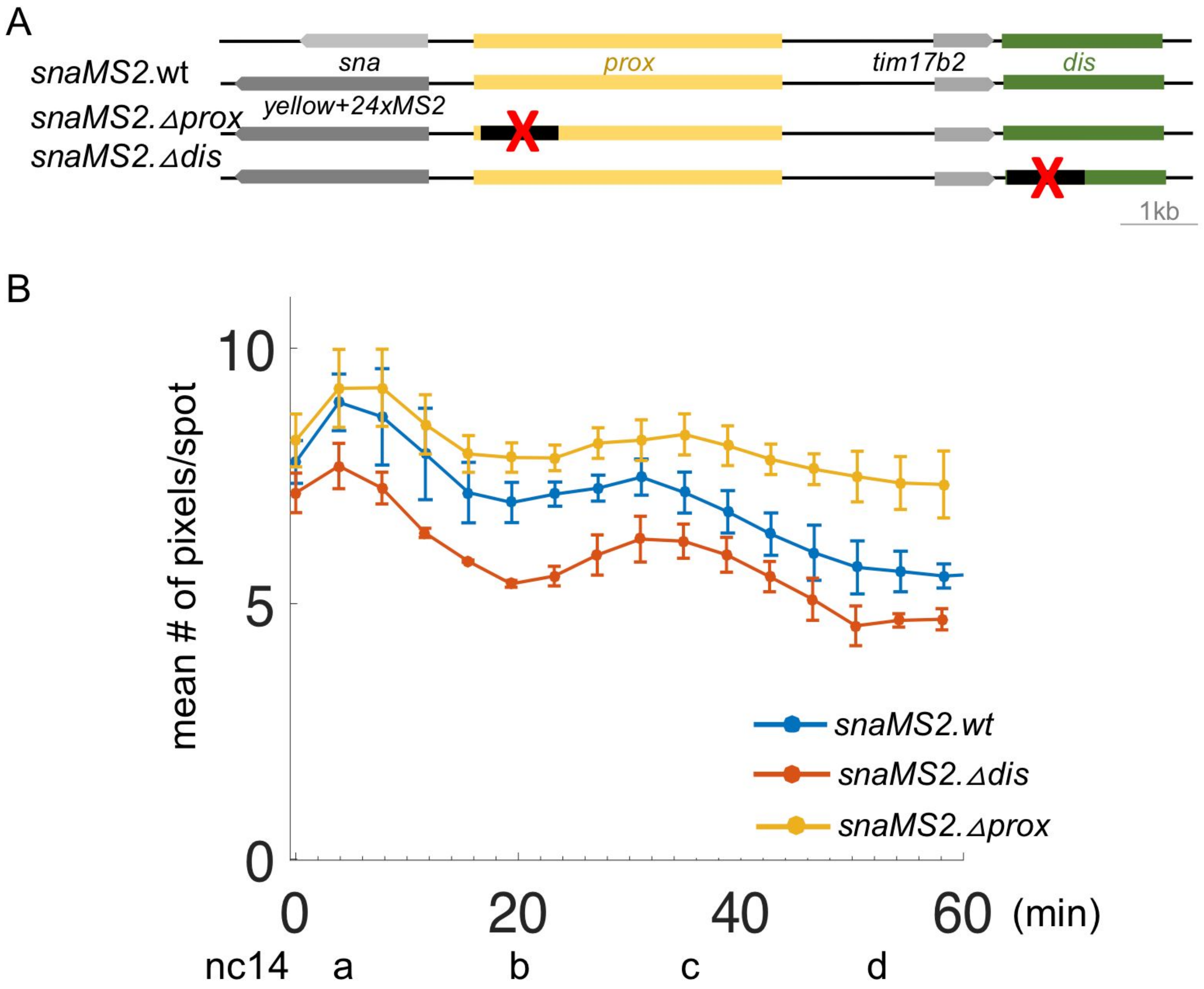

**Fig. S2. Inputs from the proximal and distal enhancers are required to support normal *sna* expression levels.** (A) Schematic of reporter constructs used to assay *sna* transcriptional activity by monitoring MS2-MCP.GFP (14). (B) Quantitative analysis of mean number of pixels per *sna* reporter MS2-MCP.GFP spot in wildtype embryos with *snaMS2.wt, snaMS2.Δprox, or snaMS2.Δdis* regulatory conditions at nc14. For each condition, movies were obtained for three individual embryos and the data averaged. Error bars represent standard error of the mean.

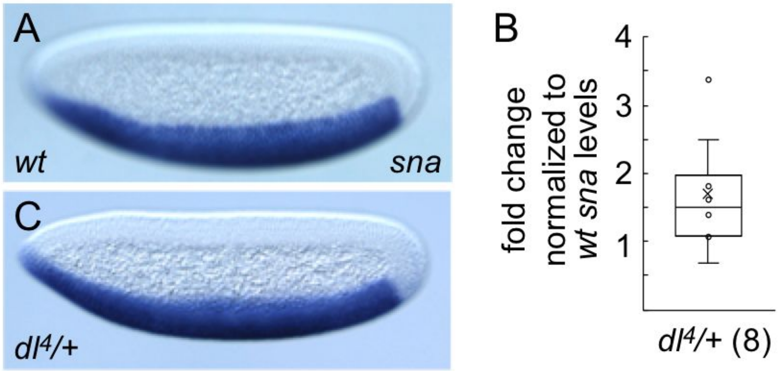

**Fig. S3. *dl*$^{4}$/+ embryos contain higher levels of *sna* transcripts than wildtype embryos.** (A,C) *sna* expression in nc14d embryos by *in situ* hybridization using *sna* riboprobes. Lateral views of embryos are shown, with anterior to the left and dorsal side up. (B) Normalized *sna* transcript levels in embryos at nc14c for indicated deletion mutations relative to wildtype determined using qPCR. Sample size n=8 embryos; also indicated next to the genotype. Box plot shows median (center line) and 25th or 75th percentiles (box limits). Whiskers represent 1.5 times the interquartile range from the 25th and 75th percentiles.

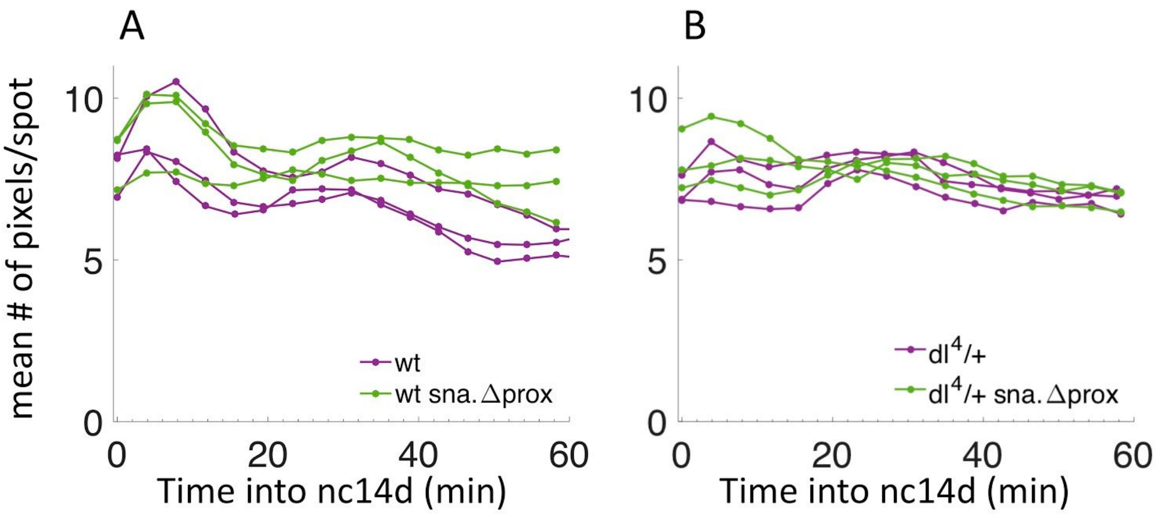

**Fig. S4. Mean number of pixels per spot associated with *snaMS2.wt* or *snaMS2.Δprox* in wildtype or *$dl^4$*/+ heterozygous individual embryos.** Each line represents the mean number of pixels per MS2-MCP.GFP spot over time for individual embryos. (A) Individual embryos plotted for a wildtype background with wt *sna* enhancers (*wt*, purple) or with the proximal enhancer deleted (*wt sna.prox*, green). (B) Individual embryos plotted for a *$dl^4$*/+ background with wt *sna* enhancers (*$dl^4$*/+, purple) or with the proximal enhancer deleted (*$dl^4$*/+ *sna.prox*, green). Some variation in starting mean number of pixels per spot is observed, but the trends remain similar. For each condition, movies were obtained for three individual embryos.

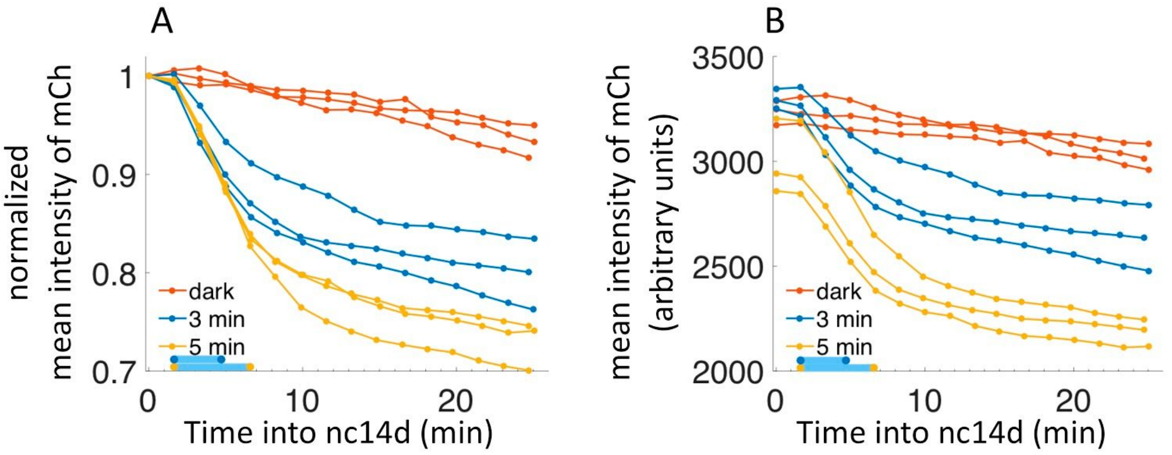

**Fig. S5. Mean intensity of mCherry fluorescence for individual embryos containing dl-mCherry-BLID with/without blue laser illumination.** (A) Plots of normalized mean intensity of mCherry for individual embryos in the dark (red), after 3 min blue laser illumination (blue), or after 5 min blue laser illumination (yellow). (B) Unnormalized plots of the same data in A. For each condition, movies were obtained for three individual embryos.

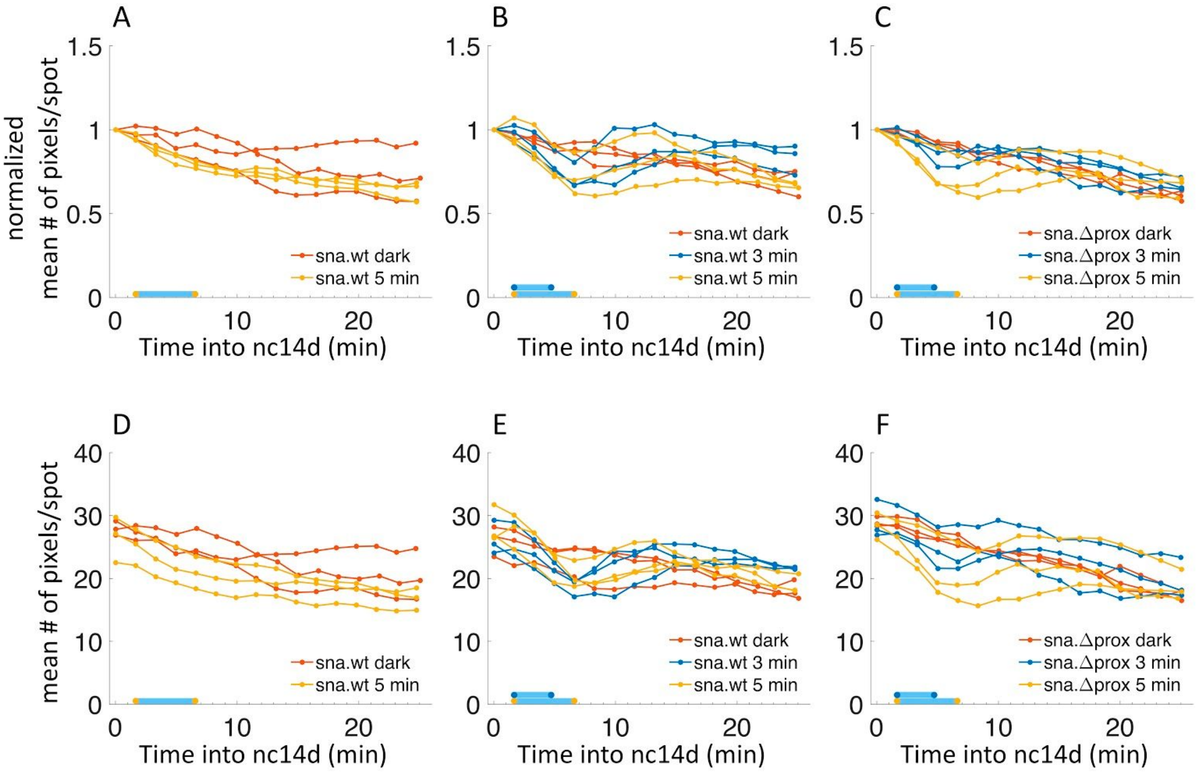

**Fig. S6 Normalized and unnormalized mean number of pixels per spot associated with *snaMS2.wt* or *snaMS2.Δprox* in wildtype or dl-BLID/+ heterozygous individual embryos under various blue laser conditions.** (A-C) Each line represents the normalized mean number of pixels per MS2-MCP.mCherry spot over time for individual embryos. (A) Individual embryos plotted for a wildtype background with wt *sna* enhancers in the dark (*sna.wt* dark, red) or with 5 min of blue laser illumination (*sna.wt* 5 min, yellow). (B) Individual embryos plotted for a *dl-BLID/*+ heterozygous background with wt *sna* enhancers in the dark (*sna.wt* dark, red), 3 min of blue laser illumination (*sna.wt* 3 min, blue), or with 5 min of blue laser illumination (*sna.wt* 5 min, yellow). (C) Individual embryos plotted for a *dl-BLID/*+ heterozygous background with the *sna* proximal enhancer in the dark (*sna.prox* dark, red), 3 min of blue laser illumination (*sna.prox* 3 min, blue), or with 5 min of blue laser illumination (*sna.prox* 5 min, yellow). (D-F) Each line represents the unnormalized mean number of pixels per spot for an individual embryo. (D) Unnormalized plots of the same data in A. (E) Unnormalized plots of the same data in B. (F)

Unnormalized plots of the same data in C. For each condition, movies were obtained for three individual embryos.

**TABLE S1. Key Resources Table**

| REAGENT or RESOURCE | SOURCE | IDENTIFIER |
|---|---|---|
| **Antibodies** | | |
| Anti-Sheep Alexa Fluor 488 | ThermoFisher | A-11015 |
| sheep anti-digoxigenin | Life Technology | PA185378 |
| **Chemicals, Peptides, and Recombinant Proteins** | | |
| Trizol Reagent | ThermoFisher | 15596018 |
| Digoxigenin labeled nucleotides | Roche | 11277073910 |
| Ex Taq DNA Polymerase | Clontech | RR001C |
| DNase I (RNase free) | NEB | M0303S |
| SuperScript III First-Strand Synthesis System | ThermoFisher | 18080051 |
| Halocarbon 27 oil | Sigma-Aldrich | MKBJ5699 |
| **Critical Commercial Assays** | | |
| SYBR Green I Master | Roche | 04707516001 |
| **Experimental Models: Organisms/Strains** | | |
| *D. melanogaster* $dl^4$/*CyO* | Bloomington *Drosophila*Stock Center, BDSC) | 7096 |
| *D. melanogaster* $dl^1$/*CyO* | BDSC | 3236 |
| *D. melanogaster dl;dl-venus* | (7) | n/a |
| *D. melanogaster nos>MCP.GFP, nos>mCherry-PCP, His2Av-eBFP2* | (22) | n/a |
| *D. melanogaster snailBAC>MS2* WT | (14) | n/a |
| *D. melanogaster snailBAC>MS2*NoPrimary | (14) | n/a |
| *D. melanogaster snailBAC>MS2*NoShadow | (14) | n/a |

| |
|---|
| *D. melanogaster y2cho2v1; P {nos-phiC31\int.NLS}6X; attP2 (III)* |
| *D. melanogaster y2cho2v1;sp/CyO;P {nos-Cas9,y+,v +} 2A* |
| *D. melanogaster* * |
| **Oligonucleotides** |
| **Recombinant DNA** |
| **Software and Algorithms** |

### Legends for Movies

**Movie S1. *snaMS2.wt* in wildtype or *dl*$^{4}$/+ heterozygotes at nc14.** Related to Figure 3C-F. MS2-MCP imaging of an embryo containing *snaMS2.wt* reporter using a 405nm laser (1.5% power) to detect H2A-BFP [i.e. His2Av.eBFP2; (22)] to identify nuclei, and a 488nm laser (2.4% laser power) to detect GFP signal associated with nascent transcripts.

**Movie S2. *snaMS2.Δprox* in wildtype or *dl*$^{4}$/+ heterozygotes at nc14.** Related to Figure 3E,F. Imaging settings are equivalent to Movie S1.

**Movie S3. Dl-mCherry-BLID imaging in late nc14 in embryos kept in dark.** Related to Figure 4B. A 561nm laser with 5% laser power was used to detect the mCherry signal associated with dl-mCherry-BLID.

**Movie S4. Dl-mCherry-BLID imaging in late nc14 in embryos with 3 min or 5 min blue laser illumination.** Related to Figure 4B. A 561nm laser with 5% laser power was used to detect the mCherry signal associated with dl-mCherry-BLID. A 488nm laser with 20% laser power was used for blue laser illumination starting at the second time frame (see Fig 4A).

**Movie S5. *snaMS2.wt* in wildtype embryos without or with 5 min blue laser illumination at late nc14.** Related to Figure 4C. Imaging settings are equivalent to Movie S3 and S4.

**Movie S6. *snaMS2.wt* in *dl-BLID/*+ heterozygous embryos without blue laser illumination at late nc14.** Related to Figure 4D,F. Imaging settings are equivalent to Movie S3.

**Movie S7. *snaMS2.wt* in *dl-BLID/+* heterozygous embryos with 3 min or 5 min blue laser illumination at late nc14.** Related to Figure 4D,G,H. Imaging settings are equivalent to Movie S4.

**Movie S8. *snaMS2.Δprox* in *dl-BLID/+* heterozygous embryos without blue laser illumination at late nc14.** Related to Figure 4E,F. Imaging settings are equivalent to Movie S3.

**Movie S9. *snaMS2.Δprox* in *dl-BLID/+* heterozygous embryos with 3 min or 5 min blue laser illumination at late nc14.** Related to Figure 4E,G,H. Imaging settings are equivalent to Movie S4.
